# Population genetic structure of disease-causing vancomycin-resistant *Enterococcus faecium* in a hospital closely reflects the dynamics of colonizing populations in patients

**DOI:** 10.64898/2026.01.09.698598

**Authors:** Camilo Barbosa, Aline Penkevich, Kevin Tracy, Steven J. Losh, Andrew F. Read, Robert J. Woods

## Abstract

Vancomycin-resistant *Enterococcus faecium* (VREfm) is a common nosocomial pathogen that can lead to severe and difficult-to-treat infections. VREfm effectively exploits healthcare environments because transmission chains are difficult to prevent and eradicate. VREfm also readily acquires antimicrobial resistance (AMR), strongly reducing the likelihood of treatment success. Here we compare the genetic population structure and antimicrobial resistance load of VREfm strains that are typically associated with patient colonization, and those found to cause disease. We use two separate but contemporaneous collections of clinical bacterial isolates in a tertiary care medical center: one consisting of *Enterococcus* from blood samples, and a second originating from a hospital wide surveillance strategy screening for VREfm in perirectal swabs of incoming patients. Using whole-genome sequencing and analysis of 2061 VREfm clones, we found that the genetic structure of colonizing and disease-causing VREfm closely resemble each other, and that sequence type 117 (ST117) played an important role in shaping these populations. The population structures of strains acquired at the hospital and those present on arrival are also remarkably similar. We further observed a high likelihood that the colonizing clone genetically resembles the disease-causing strain in patients who are both colonized and infected. Finally, the AMR gene load and distribution did not vary significantly between the blood isolates and the gut-associated strains. Altogether, our results highlight the similarities between colonizing and infectious VREfm populations and further emphasize the need to focus infection prevention strategies to minimize gut colonization.

**Author Summary:** Vancomycin-resistant *Enterococcus faecium* (VREfm) often lives harmlessly in the human gut but can cause life-threatening infections when it enters the bloodstream. Hospitals struggle with is pathogen because it spreads easily and tends to resist many antibiotics. We compared bacteria collected from patients’ guts and from their blood to ask whether the strains that cause infection are different from those that only colonize. Studying more than 2,000 bacterial genomes, we found that the two groups look remarkably similar: the same genetic types dominate in both, patients are usually infected by the same strain already living in their gut, an both groups also carried similar sets of resistance genes. Overall, this is important because it suggests that preventing colonization—not just treating infection—may therefore be the most effective way to reduce VREfm disease in hospitals.

## Introduction

Healthcare environments are strongly associated with the onset and spread of antimicrobial resistant (AMR) pathogens because antibiotic use is common (1–7). In particular, immunocompromised and chronically ill patients who often require prolonged hospital stays and intensive antibiotic exposure are at a higher risk of acquiring infections with AMR pathogens which are recalcitrant to antimicrobial treatment and often lead to treatment failure (8,9). Around 30% of healthcare acquired infections are caused by *Enterococci* (10), with those caused by vancomycin-resistant *Enterococcus faecium* (VREfm) classified by the CDC as a serious threat causing over 5000 deaths and $500M in healthcare costs in the US in 2023 (10–12). Recent studies show a strong association between *E. faecium* and health associated environments, especially among strains belonging to clade-A1, which is almost exclusively observed among hospital environments (13–16). This, together with the evolution of resistance against multiple drug classes in VREfm, limits the availability of effective treatments and highlights the need to better understand the genetic structure of this pathogen within health associated environments and the potential load of AMR genes they might be equipped with (10,17–19).

VREfm is a human gut commensal, but can cause severe disease as blood stream or soft-wound infections (20). Colonized patients, as well as contaminated surfaces, can facilitate the transmission of VREfm from patient-to-patient or from personnel-to-patient thus sustaining repeated transmission events and routes (13–15,21). An understanding of transmission routes can inform strategies to limit VREfm acquisition within hospital environments. So too could an understanding of how commensal VREfm transition to causing disease. Gastrointestinal tract colonization with VREfm is a substantial risk factor for the development of VREfm bacteremia (15,22–33). Moreover, whole-genome sequencing of clinical isolates have shown that cancer, hematology, and immunocompromised patients are more likely to acquire VREfm infections with strains genetically close to those found in their own gastrointestinal tracts (15,16,22,23). The implication from these studies is that a patient’s commensal VREfm population is the source of their infections. If so, preventing acquisition of commensal VREfm would reduce infection risk.

However, to date genomic surveys have been limited to less than 400 *E. faecium* genomes from at most ∼100 patients, collected for the most part over less than 6 months (15,16,22,23; Chilambi et al. (16) collected over 10y). Importantly, genomic studies evaluating blood and perianal swab isolates have been limited to 24 patients only (16) Here we substantially increase the depth and breadth of the genomic samples with 3,400 clones from almost 2,000 patients (34), including >100 patients from which we have instances of both colonization and bloodstream infection with VREfm.

We analyzed 2061 VREfm genomes from 1957 patients. These came from a collection obtained from patients at the University of Michigan Hospital between 2013-2021, comprising more than 75000 tests from over 39000 patients. We asked if, (1) the population genomic structure of VREfm from patients with blood stream infections (BL) differed from those derived from a larger population associated with colonization events as determined from perirectal (PR) swabs, (2) whether the population genomic structure of strains introduced into the hospital was different from that of strains more likely to have been acquired in the hospital environment, (3) whether the sequence type at different body sites (blood and rectum) from the same patient were different, and finally, (4) whether the AMR gene load differed significantly between the BL and PR populations. In all four comparisons, the differences were minimal. All VREfm genomes were previously reported and analyzed by our group evaluating bacteriocin dynamics in VREfm in Garretto et al., 2024 (34).

## Results

### Data structure and group composition

Our datasets were derived from two large collections from the University of Michigan Hospital (Fig 1). The first was obtained from perirectal (PR) swabs from patients as part of a hospital-wide surveillance effort screening patients for the presence of VREfm in the gut between January 1^st^, 2016, and December 31^st^, 2020. We evaluated 55666 tests for VREfm from 28878 patients. Between 5-10% of all tests were positive for VREfm during the 5y period of our study, with prevalence of VREfm decreasing over time (S1 Fig). We have stored over 5000 VREfm clones from these patients (Fig 1).

**Figure 1.**
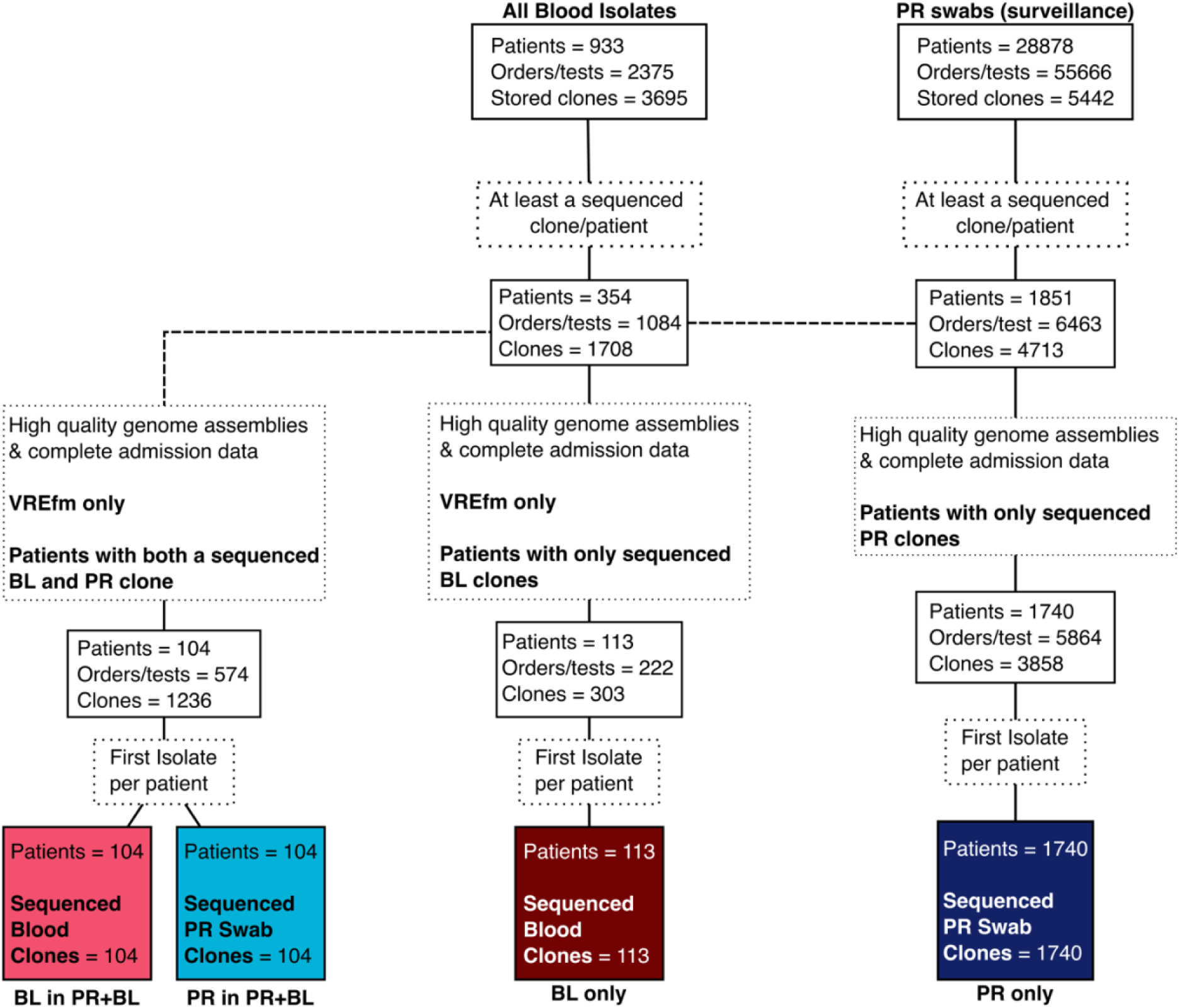
Data structure and clone selection. Isolates were obtained from two large collections. These collections were generated at the University of Michigan Hospital from 2013 to 2020. One of the collections included all blood isolates (BL) positive for *Enterococcus*, while the second collection includes isolates obtained from a perirectal swab (PR) surveillance effort screening for VREfm at various wards within the hospital from 2016 to 2020. From these collections we then selected the first sequenced clone from each patient and obtained high quality genome assemblies of VREfm for 2061 clones from 1957 patients distributed among 4 distinct groups (from left to right): 1) Patients who concomitantly had a sequenced blood clone (BL in PR+BL, pink) and a 2) sequenced PR swab clone (PR in PR+BL, light blue), 3) patients who only had BL sequenced clones (BL only, dark red), and 4) patients who only had PR sequenced clones (PR only, dark blue).

The second collection consists of bacteremia samples (BL) positive for *Enterococcus* haphazardly collected between 2013-2016 but predominantly comprised of *Enterococcus spp* obtained between 2016 and 2021 (Fig 1). These came from a total of 933 patients and 2375 test orders, resulting in 3695 enterococcal clones (see S2 Fig for a breakdown of total clone number and changes of clone numbers per species over time within the BL group).

In this study, we focus on clones confirmed as VREfm by the microbiology lab at the University of Michigan Hospital. For a subset of all obtained clones, we performed whole-genome sequencing (WGS) and generated *de novo* genome assemblies for at least one clone from 2530 patients. We then filtered high-quality genome assemblies having more than 30x coverage, 95% mapping and a minimum N50 larger than 1000 (Fig 1, and S3 Fig). Finally, we included sequenced clones for which we had complete patient’s admission and discharge data (Fig 1). We found no discernable difference in genome length or GC content between the four main groups (S4 Fig). All included genomes were previously reported and analyzed in Garretto et al., 2024 (34).

We then wanted to distinguish between patients who had (1) both a BL and PR clone during their admission (i.e., patients colonized and infected, hereafter denominated PR+BL), (2) patients who only had identified BL clones and no PR clones (hereafter denominated BL only), and (3) patients who only had PR clones and no BL clones (hereafter denominated PR only). In the PR+BL group, we identified 104 patients who had at least one sequenced BL and PR clone. In the BL only group, we found 113 patients with at least a sequenced BL clone, and no PR clones (Fig 1). For the PR only group, we obtained a total of 1740 patients with at least one sequenced clone, and a total of 3858 clones. Finally, for downstream analysis, we included only the first sequenced clone of each patient, resulting in 2061 sequenced clones obtained from 1957 patients distributed among 4 groups (Fig 1): 1) BL clones in PR+BL, 2) PR clones in PR+BL, 3) BL clones only, and 4) PR clones only.

Given *E. faecium*’s strong association with healthcare environments we wanted to determine the proportion of patients for which colonization and/or infection was suspected to have occurred within the hospital (Table 1). For the BL only, and BL in PR+BL, we followed the CDC and NHSN criteria (35) defining an infection occurring after the third calendar day of admission to an inpatient location as a hospital-acquired infection (HAI). Furthermore, our PR dataset includes weekly surveillance data whereby patients can be positive for VREfm on arrival (within 72 h of admission), positive after 72 h of admission without a previous negative swab (late positive) or have negative swabs before having a positive swab. Based on this, we further defined for the PR dataset patients with a negative swab followed by a positive one as hospital acquired. Finally, as patients could have more than one admission with multiple PR and/or BL clones, we focused on the admission from which the clones (Fig. 1) were sequenced.

**Table 1.** Proportion of colonization and infection within and outside the hospital.

| Main group | Positive on arrival | Late Positive | Hospital acquired |
| --- | --- | --- | --- |
| **PR only** | 1036/1740 (59.5%) | 205/1740 (11.8%) | 499/1740 (28.7%) |
| **PR in PR+BL** | 51/104 (49%) | 8/104 (7.7%) | 45/104 (43.3%) |
| **BL only** | 53/113 (46.9%) | NA | 60/113 (53.1%) |
| **BL in PR+BL** | 30/104 (28.8%) | NA | 74/104 (71.2%) |

Within the PR groups (PR only and PR in PR+BL), we found that patients that were both colonized and had bacteremia more readily acquire the PR clone at the hospital than those patients who are only colonized (Χ^2^ = 7.95, P = 0.019; Table 1). This was also observed for the BL clone in patients both colonized and infected (Χ^2^ = 19.31, P < 0.001).

### Bacteremia associated VREfm from before 2016 have a distinct genetic structure

We used multi-locus sequence typing (ST) to identify differences in the population structure between our four main groups in a broad sense. We found a total of 37 different STs across all groups (Fig 2A). Patients colonized with VREfm but without bacteremia (PR only), had a higher richness (35 different STs) than patients in other groups, due to greater sample size (Fig 2A). However, despite being the richest group, patients with bacteremia had have similar diversity though BL only is slightly more diverse (rarefaction curves in Fig 2B, and Simpson’s diversity in S5 Fig). ST1703 was exclusively observed among the BL and PR clones in patients both colonized and infected (PR+BL), with all other STs in these groups being shared with the PR and BL only groups (Fig 2C). Notably, the 22 STs exclusive to the PR only group, account for less than 10% of the total population (Fig 2C and 3A).

**Fig. 2.**
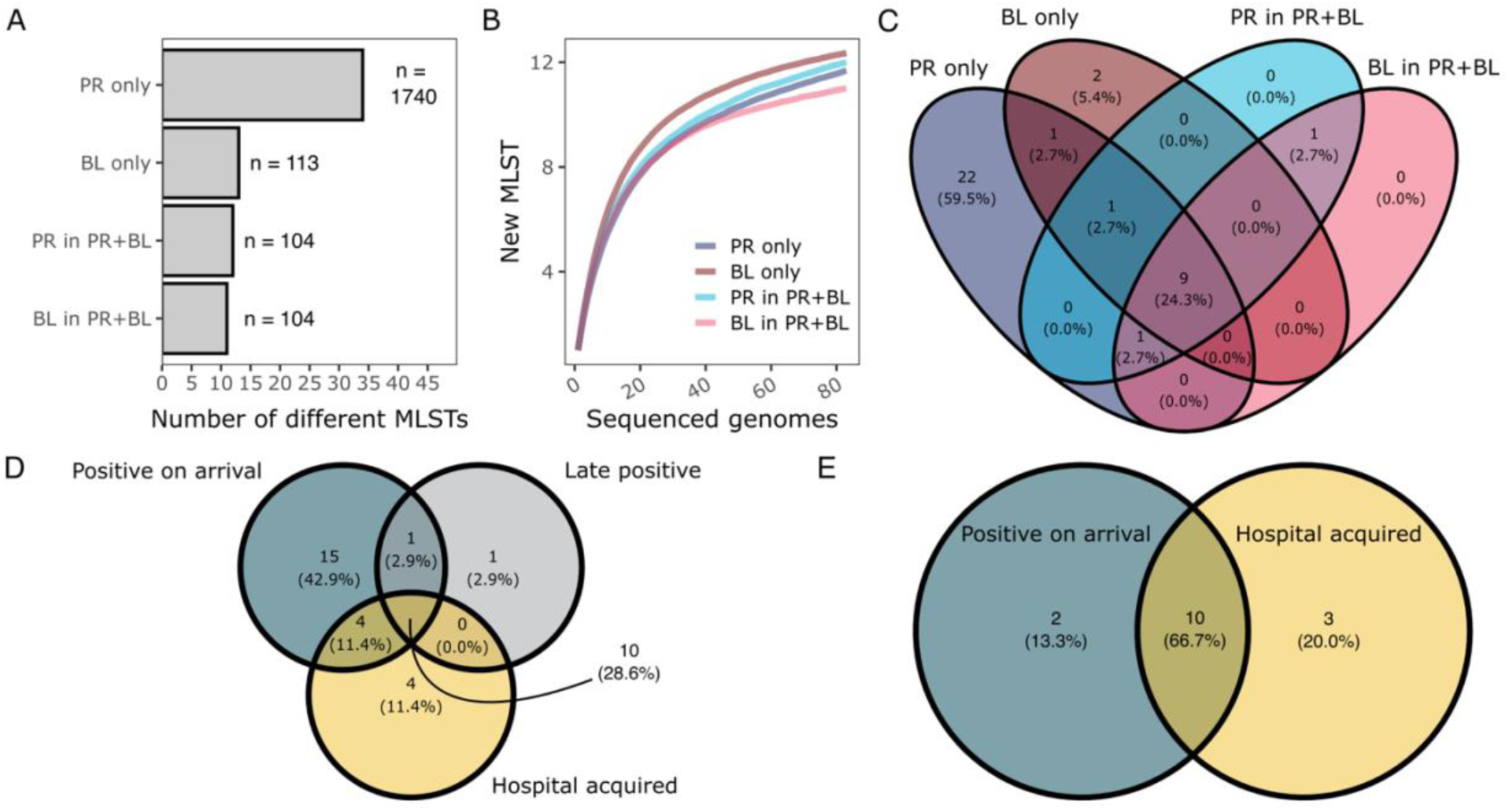
MLST diversity and richness between groups. In (A), total richness of MLSTs is shown for each group as well as the total number of patients/sequenced clone (n). Diversity was calculated using rarefaction curve analysis for all groups using the lowest number of patients/sequenced clones. Curves (B) correspond to the mean of 1000 sampling iterations per group. Venn diagrams of shared MLSTs between groups (C), pooled PR clones analyzed by time of colonization/infection (D), and pooled BL clones analyzed by time of infection (E).

**Figure 3.**
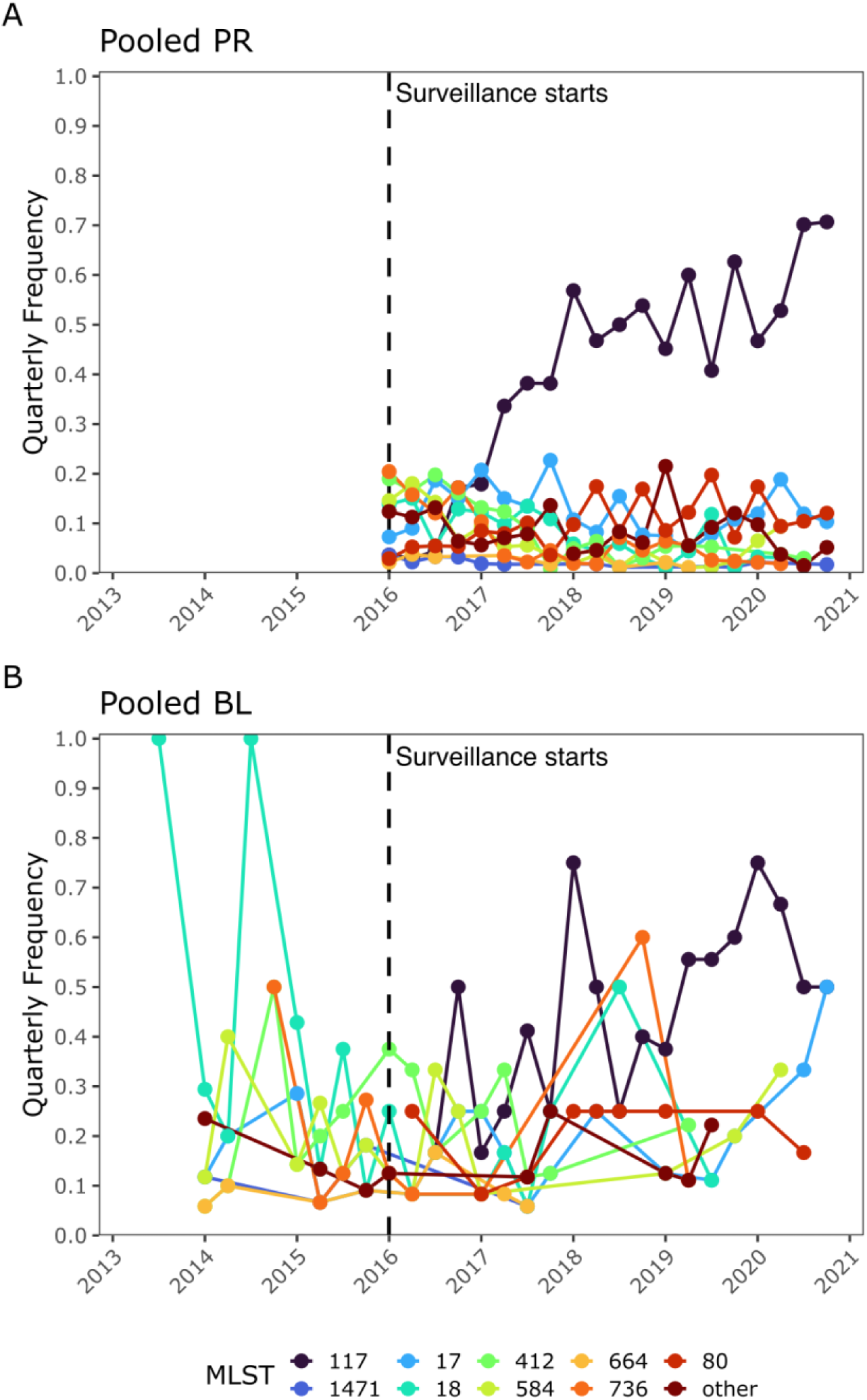
Changes in MLST frequency over time. Quarterly frequencies for the 10 most common MLSTs in the pooled PR clones (A), and pooled BL clones (B). The PR surveillance was initiated in 2016, but BL clones were obtained before this time. The vertical dashed line indicates the beginning of the surveillance strategy at the University of Michigan Hospital. Fig 3A is partly reconstructed from Fig 8B in Garretto et al., 2024 (34).

To determine if the clones seen on arrival were genetically distinct from all others, we pooled all PR clones together (Fig 2D), as well as all BL clones (Fig 2E), and analyzed them relative to their time of colonization. For the pooled PR group (Fig 2D) we found that around 43% of the STs were exclusively observed among clones from patients positive on arrival (among 1087/1844) clones; see Table 1). In contrast, only four (11.4%) STs were found exclusively among PR clones acquired at the hospital (among 551/1860 clones; see Table 1). Ten STs were observed in all colonization time groups among all the pooled PR clones. Among the pooled BL clones, we found a similar pattern whereby 66.7% of all STs were predominantly observed among patients positive on arrival, with three STs being exclusive to HAI among 83/217 BL clones (Fig 2E and Table 1). Interestingly, the STs unique to HAI are different among the PR and BL clones: STs 64, 94, 666 and 1478 were only observed among PR clones, and STs 1516, 1660 and 203 were exclusively seen among BL clones. These latter sequence types are however among the least frequently observed ones across all different groups (Fig 4).

**Figure 4.**
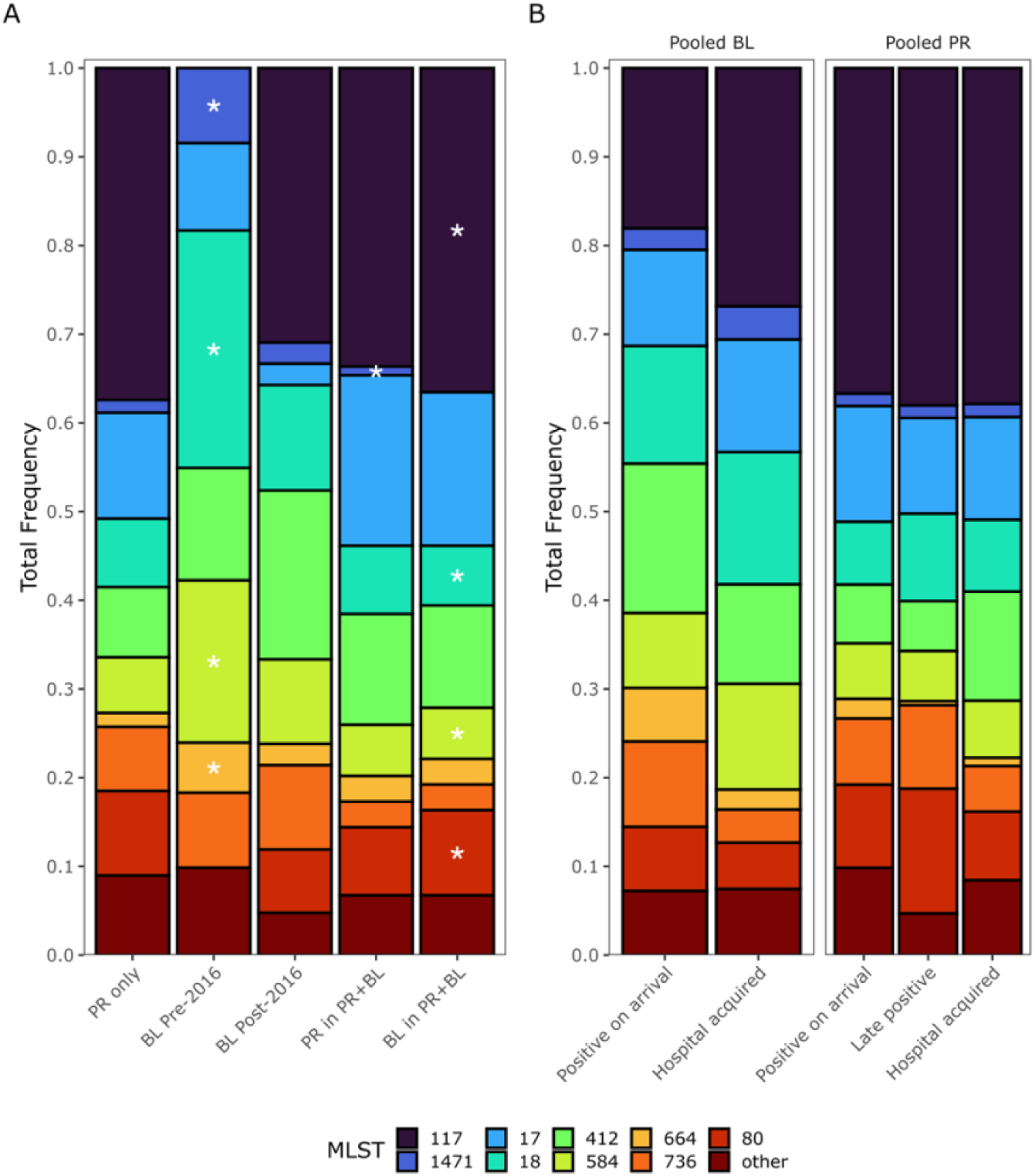
Total ST frequency. The total frequencies of the top 10 STs were calculated for each group (A), and for colonization time (B) among the pooled BL (right panel) and the pooled PR (left panel). We asked if the frequencies observed for the BL only before and after the surveillance started, and the BL in PR+BL, were a random subsample of the pooled PR frequencies (PR only and PR in PR+BL) in a randomization test. Low frequency STs and genomes with no assigned ST were grouped as “other”. Frequencies with a significantly different frequency for each ST and group are highlighted with a white asterisk (*), In (A), STs 117 and 80 in the BL pre-2016 had a frequency of zero and were significantly different in the randomization test, as well as ST 1471 in the BL in PR+BL group. P-values were adjusted for multiple testing using the FDR method (36).

To assess statistical significance, we focus on the frequencies (and absolute numbers) in the 10 most common ST changed over time (Fig 3 and S6 Fig). Among the PR clones, we observed that ST117 has become predominant since its first appearance in the first quarter of 2016 (Fig 3A and S7 Fig). ST80 is also steadily increasing in frequency over time, albeit at a much lower rate than ST117. Similarly, ST117 and ST80 have increased consistently in frequency among the BL clones after they were first observed in the third and second quarter of 2016, respectively. Importantly, neither of these STs were observed before this point in time among the BL clones, thus suggesting that their dominance within this hospital is a recent trend that has shaped the genetic structures of both the BL and the PR populations. Among the BL population, ST117 and ST80 have displaced ST18 which was the most predominant genotype before 2016.

We then analyzed the population structure of the PR and BL groups by considering the total frequency of the top 10 STs (Fig 3). We used a randomization test with replacement to determine if the observed distributions of STs in the BL only and BL in PR+BL groups were consistent with subsamples of the pooled distribution of the pooled PR clones in PR and PR in PR+BL. Since the BL population included clones before 2016 which marked the beginning of the PR surveillance strategy, we analyzed the population structure of the BL group before and after 2016. Briefly, we sampled from the pooled PR clone’s distribution the corresponding sample size of each of the BL groups and calculated the frequencies of the observed STs. We repeated this process 10,000 times, calculated the probability of obtaining the observed frequency of each group for each ST and adjusted significance for multiple testing using the False-discovery rate (FDR) method (36).

Overall, we found that the BL only population from before 2016 and the BL in PR+BL were significantly different from the overall PR population distribution while all other were statistically indistinguishable from it. This is also consistent with the described temporal trends, whereas all other groups closely resembled it (Fig 3A and 4A). In the BL only population 6/10 STs (Before 2016, ST117 and ST80 were not observed and thus had a frequency of 0, and were statistically distinct from the observed frequencies in the PR population) had a significantly different distribution than expected by chance (Fig 4A) while 5/10 where significantly different in the BL in PR+BL group (ST1471 had a frequency of 0 and was significantly different as well, but not visible in Fig 4A). Importantly, before 2016, ST117 and ST80 were not observed but had a similar distribution as in the PR population after this time, again highlighting their dominance within this hospital. Furthermore, the population structures of BL clones after 2016 is remarkably like that of the pooled PR clones. The lack of PR surveillance data before 2016 does not allow us to determine whether the VREfm population associated with bacteremia is a continuous subpopulation of that of VREfm associated with gut colonization as observed after 2016. However, our data strongly emphasizes the impact the introduction of ST117 and ST80 have had on both populations. Overall, we found that that pre-2016 the BL population has a different ST distribution than the PR population, whilst all other groups are remarkably similar.

We further wanted to assess whether there were differences in the population’s structure when considering the time of colonization/infection (Fig 4B). For this we pooled the BL and PR clones and for each we then looked at the frequency of each of the top STs relative to the time of colonization/infection (Fig 4B). To determine differences in the colonization time we did a similar analysis as above where we determined if the distribution of STs represented a random subsample of the pooled PR or BL population structures, respectively. In this case we used pooled BL and PR population structures as reference distributions for their corresponding groups (Fig 4B). After correcting for multiple testing, we found no significant differences in the frequency of hospital acquired or incoming clones relative to the pooled PR distribution, thus suggesting a general common population structure for clones of patients colonized/infected on arrival and that of patients getting colonized/infected in the hospital (Fig 4B).

### Patients have a high likelihood of being colonized and infected by the same ST

Considering the previous observation that the distribution of BL clones in the PR+BL group resembles that of the PR only group, and that 88/104 patients had a matching STs between the BL and PR pairs in each patient, we wanted to determine the probability of the STs in the BL and PR pairs in the PR+BL groups to match as a function of time (Fig 5 and S8 Fig). To approach this, we employed a logistic regression as a function of time between the two clones for three distinct cases within the PR+BL group: 1) in all PR+BL patients, using the absolute value of the time between the clones, 2) patients in which the PR clone was observed before the BL clone (indicative of colonization preceding infection), and 3) patients in which the BL clone was observed first (Fig. 5A-C). For the first case, we found that if the clones were sampled on the same day, the predicted probability of them being the same ST was 0.93 (95% CI = 0.97 - 0.85, Fig. 5A). That probability dropped the longer the time between clone isolation, with the uncertainty around the probability of a match increasing as well. Importantly, this pattern holds regardless of whether the BL or the PR clone was found first (Fig. 5B and 5C). Similarly, we found that the probability of having a positive PR clone for VREfm is roughly the same shortly before or after identification of the BL clone, with the probability decreasing over time if the BL clone is isolated before the PR clone, but not on the reverse case even if the clones are 90 days apart from each other (S9 Fig). Overall, this demonstrates that when the colonizing population is sampled close in time to the blood isolate, the likelihood of recovering the same ST is considerable.

**Figure 5.**
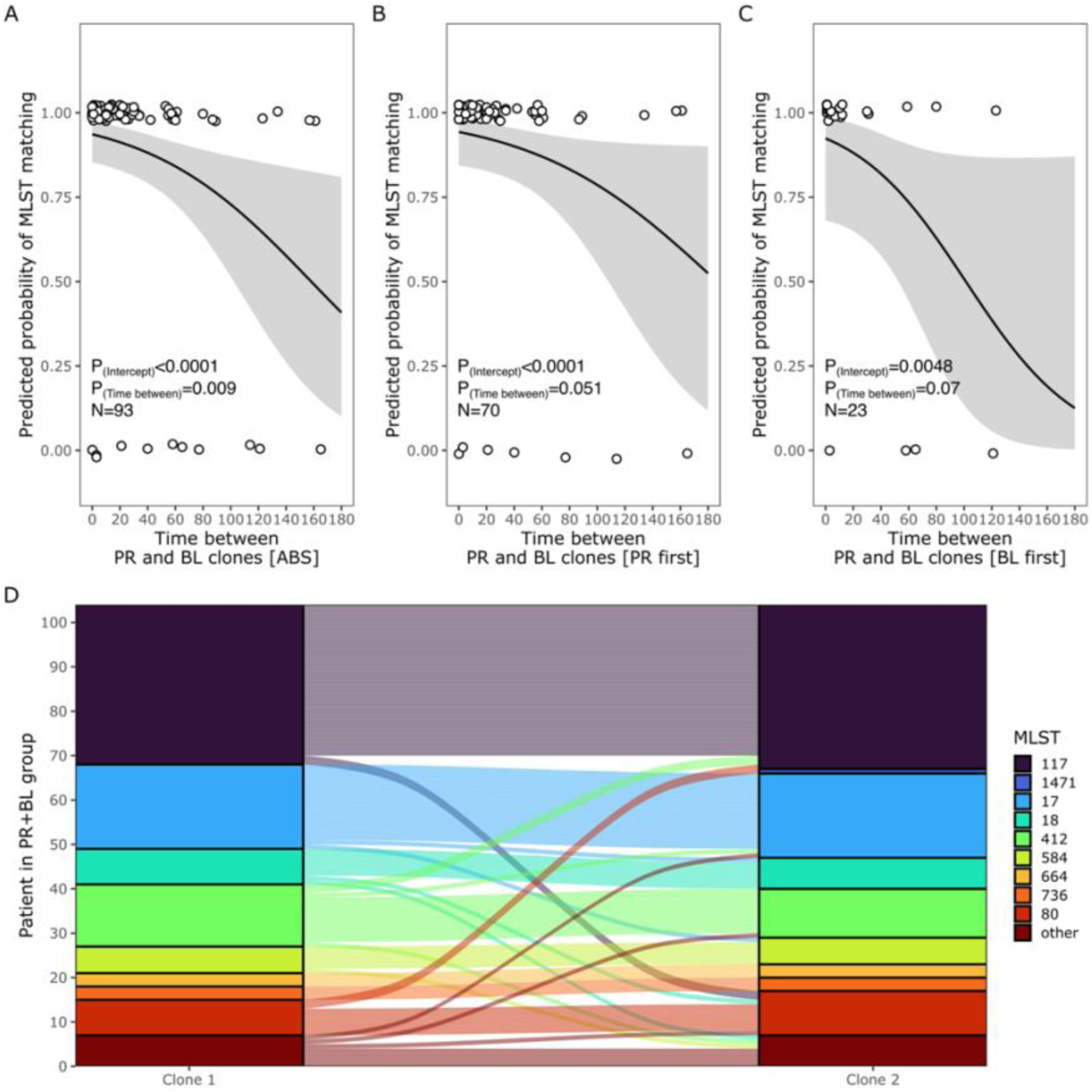
Probability of isolating PR and BL clones with matching ST in the PR+BL group as a function of time of isolation between the clones. We used logistic regression to calculate the predicted probability of ST matching as a function of time regardless of which isolate was identified first (A), or by assuming that colonization (PR clone obtained first, B) precedes the BL clone, or that it does not (C). (D) Alluvial plot of ST mismatch between the first and second clones in patients with both PR and BL clones (PR+BL).

In the 23 cases where we found a mismatch between the STs, we observed some interesting patterns (Fig. 5D and Fig. S8): First, when the first clone is ST117 and there is a mismatch, the second clone is exclusively ST80. Second, when there is a mismatch and the first clone’s genotype is ST412, the second clone is ST117. Interestingly, ST117 is only mismatched by ST412. Third, the most common ST of the second clone when there is a mismatch is either ST80 or the low frequency STs grouped in ‘other’. Finally, ST664 and ST736 are the only genotypes that always match the first and second clones.

Based on a previous study examining for the presence of bacteriocin Bac43 among our isolates (34), we additionally considered how often the second isolate in PR+BL group of isolates acquired this bacteriocin as an indication of a biological advantage to colonize/infect hosts. Among the first isolate we found that half (52) of the clones had Bac43, while 60 of the second isolate carried it. This increase was not statistically significant (Χ^2^ = 0.95, P = 0.33, See Fig S10)

### AMR genes/variants are equally distributed in bacteremia associated pathogens and colonizing strains

To determine the AMR gene load in our clones, we used AMRFinder to identify genes and variants commonly associated to antibiotic resistance against multiple antibiotics in *E. faecium*. From this, we identified a total of 61 distinct genes and/or variants associated to AMR, predominantly seen in the PR only group which was the richest (Fig 6A). Despite the difference in numbers, and PR only being the richest group in terms of AMR genes/variants, all groups had similar levels of diversity (Fig 6B and S11 Fig). In general, we observed a similar pattern as in our ST analysis, with the much larger PR only group having many AMR genes/variants observed only in this group. However, around 50% of all the identified genes/variants were shared among all groups, a higher proportion than that of shared STs (Fig 6C and Fig 2C). Similarly, when we analyzed differences between clones from patients positive on arrival, late positive and hospital-acquired, we found that 62% of all the AMR genes/variants were shared among PR clones (Fig 6D), and more than 80% among the BL clones (Fig 6E). At the clone level, we found no significant differences in the number of AMR genes/variants carried by the clones in the different groups (Fig 6F, P = 0.0676), with a median of 19 AMR genes/variants. However, we found significant differences in the number of AMR genes/variants between STs (Fig 6G and S1 Table). In general, ST117, ST17, ST18 and ST412 were significantly different (S1 Table) to most other MLSTs having a lower number of AMR genes/variants (Fig 6G).

**Figure 6.**
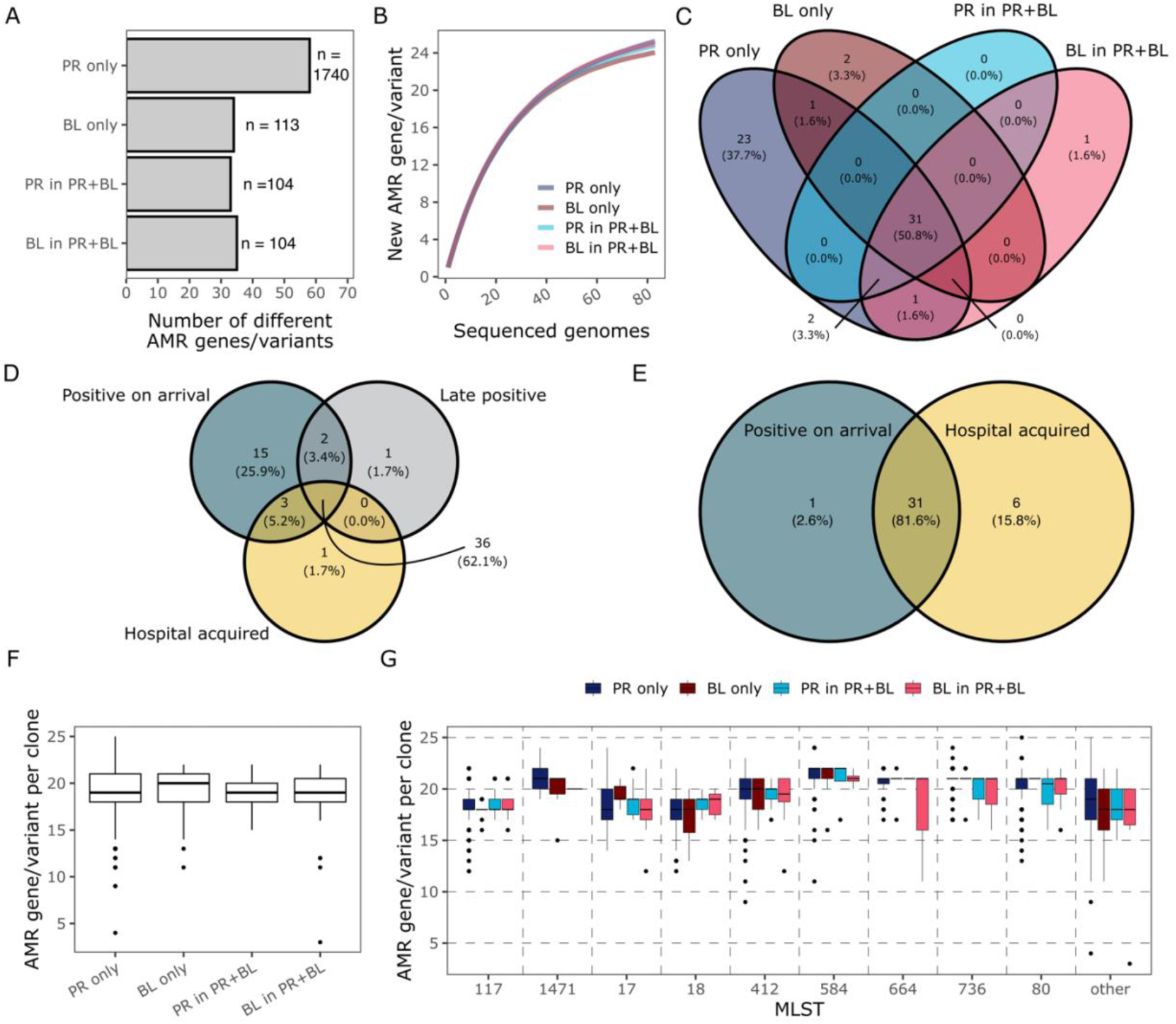
Richness and diversity of AMR associated genes and variants among the different groups. Genes associated to antibiotic resistance in *E. faecium* were identified using AMRFinder. The total number of distinct AMR genes and variants were calculated for each group (A). Diversity was calculated using rarefaction curve analysis for all groups using the lowest number of patients/sequenced clones. Curves (B) correspond to the mean of 1000 sampling iterations per group. Venn diagrams of shared AMR genes/variants between groups (C), pooled PR clones analyzed by time of colonization/infection (D), and pooled BL clones analyzed by time of infection (E). We calculated the number of AMR genes and variants per clone for each group (F) and for the top STs (G). An additive two-way ANOVA using the Type III sums of squares method to account for unbalanced groups was used to identify significant differences in the number of AMR genes/variants per clone between the 4 groups, and STs. We did not find significant differences between the 4 main groups (P = 0.068) but found significant differences between STs (S1 Table).

We then asked how similar the AMR gene load was between BL and PR clones, and between hospital acquired and positive on arrival patients (Fig 7). We calculated the frequency of each gene for the pooled PR clones and compared it to that of the clones in the BL only (Fig 7A) and the BL in PR+BL (Fig 7B). We expected the load of AMR genes/variants to be the same between these two groups and thus align close to a 1:1 slope line (black line ± 95% CI dashed lines). Indeed, most of the identified AMR genes/variants had a similar frequency in the pooled PR and BL only clones, with all genes having a similar frequency between the pooled PR and the BL in PR+BL clones (Fig 7A and B). In both cases, the slope from a linear regression was close to one and R^2^ above 0.94 (m_BL_ = 0.952, m_BL in PR+BL_ = 1.02 and R^2^_BL_ = 0.94, R^2^_BL in PR+BL_ = 0.995 for the BL only and the BL in PR+BL groups, respectively; Fig 7A and B). The few genes with dissimilar frequencies between the BL only clones and the pooled PR clones included *parC* and *parR*, which are typically associated with quinolone resistance in various species (37,38), as well as *dfrF* and *aac(6’)-Ie*-aph(2’’)-Ia which are associated to resistance against trimethoprim and aminoglycosides, respectively (39–41). Overall, this suggests that, with a few exceptions, the AMR genes/variants present in isolates associated to bacteremia and colonization is remarkably similar. We had the same expectation when comparing the frequencies based on time of colonization for the pooled PR and BL clones (positive on arrival vs. hospital acquired; Fig 7C and D). The frequency between the two groups is almost identical (respectively, Fig 7C and D).

**Figure 7.**
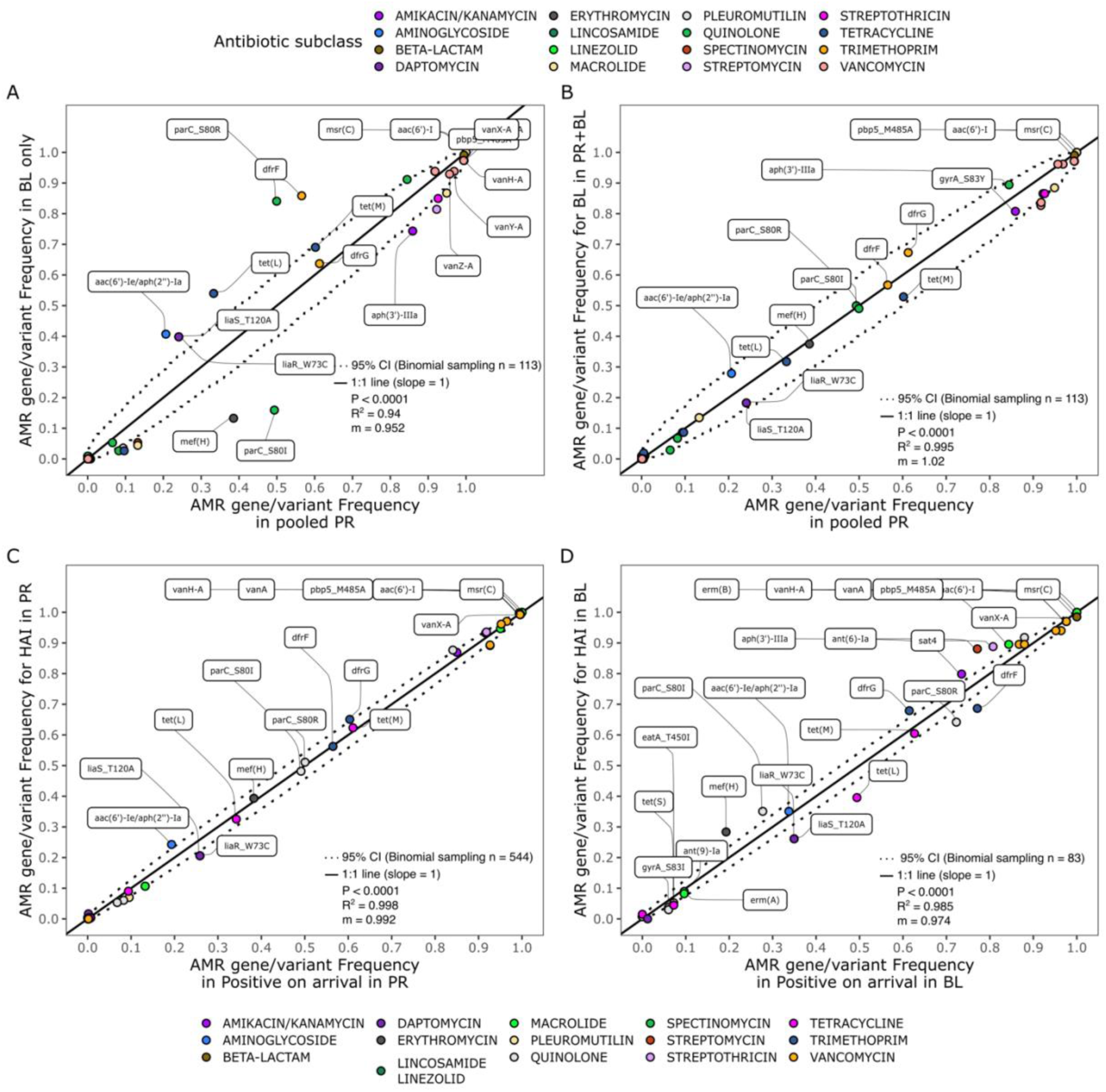
The frequencies of AMR genes and variants in blood isolate clones form patients who were also colonized differs from those who did not present a PR clone. Frequencies for the 61 AMR genes/variants identified across all sequenced clones were pooled for all PR clones and analyzed relative to those of BL only sequenced clones (A) and the BL clones in PR+BL (B). Similarly, we analyzed the frequencies of all genes within the clones acquired at the hospital (HAI) relative to those positive on arrival for the pooled PR (C) and BL (D) clones. A linear regression was calculated for each group (dashed line and grey area indicate the regression line ± se), with P values and R^2^ indicated within the corresponding panel. A 1:1 line (slope =1) is also shown (black). Each point corresponds to an AMR gene or variant, with the different colors highlighting the antibiotic subclass they are associated with.

## Discussion

Our underlying hypothesis was that the genetic population structure of clones associated with bacteremia is tightly associated with that of the clones linked with gut colonization. This is important because if true, prevention and treatment strategies that minimize colonization and subsequent transition from the gut to other body sites would minimize the risk of infection regardless of the ST or the population structure of the colonizing population. We asked four questions. First, did the population genomic structure of VREfm from patients with blood stream infections (BL) differ from those derived from a larger population associated with colonization events as determined from perirectal (PR) swabs? We found that it did not: the genetic population structure of bacteremia-associated clones changed dramatically over time, driven largely by the spread of ST117, but these temporal changes tightly resemble the population distribution of colonizing clones (Fig 2-4). Second, did the population genetic structure of strains introduced into the hospital differ from that of strains more likely to have been acquired in the hospital environment? We found that it did not (Fig 3, 4). Third, were the sequence types associated with colonization and septicemia from the same patient congruent? We found a high likelihood that they were (Fig 5). And finally, did the AMR gene load differed significantly between the commensal and septicemic populations differ. We found the differences were minimal (Fig 6 and 7).

These results are consistent with previous findings suggesting that gut colonization is a significant risk factor for the development of bacteremia (15,16,22,23). For instance, when the PR and BL clones were detected contemporaneously or nearly so, they were highly likely to be of the same sequence type, but the greater the time between detection of a PR clone and a BL clone, the less likely they were to be of the same sequence type (Fig 5A-C), suggesting it is not uncommon for patient to harbor multiple VREfm. Previous studies have shown that gut colonization typically precedes bloodstream infection, particularly in vulnerable patients (24,30,33). Formally, directionality (gut to bloodstream versus bloodstream to gut) is hard to determine, though we found VREfm was more likely to be detected in a patient’s PR swab before VREfm septicemia rather than after (Fig 5).

Our dataset allowed us to potentially distinguish between patients who were colonized or infected on arrival or acquired it at the hospital. Overall, we found that in cases of patients both colonized and infected, there was a higher likelihood that both the PR and BL clone were acquired at the hospital. This is not surprising as prolonged hospitalization is a risk factor both VREfm colonization and infection (22,24,42). Moreover, our genomic analysis emphasized the remarkable similarities between the population structures of clones present on patient arrival and those infected at the hospital. Interestingly, we observed that around 70% of colonized patients arrive at the hospital already carrying VREfm, suggesting the pattern in this hospital may reflect processes in the broader community and healthcare system (43). In sum, this further suggests that efforts to curve the prevalence of infections with VREfm need to center on preventing any VREfm colonization, without focusing on specific STs.

We did not find a significant difference between the amount or frequency of AMR genes between clones associated with colonization and those associated with bacteremia. This was particularly true for clones obtained from patients having both a PR and BL clones (PR+BL), where the frequency of all genes was almost in a 1:1 ratio, as well as those from patients positive on arrival and those acquired in the hospital. For patients with only BL clones identified, the relationship was not as strong, with less than 10 genes having disparate proportions between the two groups. Importantly, most of these genes are associated with resistance against antibiotics that are not typically used to treat VREfm, for which linezolid and daptomycin are typically recommended (44,45), and commonly used within this hospital (46). We found that 40% of the BL isolates had close to 1.6 times more mutations in the three-component regulatory system *liaSR* which is linked to the cell envelope stress response that is also associated with varying levels of resistance against daptomycin (41,47). This difference could be associated with subtle difference in the ST distribution in the BL only group, as a recent study found a strong association between these mutations and a variety of STs (48). We observed a similar pattern for *aac(6’)-Ie-aph(2’’)-Ia* which encodes for the most common aminoglycoside modifying enzyme among enterococci (39,49). Due to their toxicity, aminoglycosides are not typically considered for treatment, except when used in combination with β-lactams (44). Ultimately, regardless of the AMR gene/variant we did not find significant differences among the main groups, thus further highlighting the genetic similarity between these two populations (BL and PR).

We also found mutations in genes associated with resistance against antibiotics more commonly used to treat other type of infections. For instance, around 80% of the BL clones, and 55% of the PR clones, had the *dfrF* gene, a dihydrofolate reductase typically associated to resistance against trimethoprim in several species (40,50). Trimethoprim is more commonly used for the treatment of enterococcal urinary tract infections, and not for bacteremia (51). This pattern may be reflective of distinct transmission chains originating in distinct wards of the hospital or the likely result of off-target antimicrobial therapy for the treatment of non-enterococcal infections.

ST117 has been reported to be an emergent nosocomial, multidrug-resistant pathogen worldwide (20,26,52–55). We first detected ST117 in among our first of our samples of PR clones in February 2016, in a patient positive on arrival. ST117 was not observed among the BL clones until after the third quarter of 2016 (even though we have some BL samples back to 2013 (Fig 3 and S4 Fig). After its appearance, ST117 steadily increased in frequency in colonized patients and in those with bacteremia, displacing the previously more common ST18, ST412 and ST584 among patients with bacteremia.

The factors that facilitate ST117’s success within hospital environments remain unclear. We found no evidence suggesting that ST117 has a significant advantage of being acquired at the hospital (Table 1), or that it was more likely to cause an infection after colonization with a different ST, or that it disproportionally harbored more AMR genes than other STs. Several studies have attempted to look for the presence of virulence factors and/or antimicrobial resistance genes that could account for ST117 predominance worldwide over the last 10 years and generally conclude that AMR genes in ST117 strains that were also readily found in other sequence types (27,52,56–67). For example, several ST117 harbor *vanB*, a vancomycin resistance gene, in the integrative conjugative element (ICE) Tn1549 often associated with gut anaerobes (27). Yet, this ICE and other variants of it and the *vanB* cassette, are also commonly observed in other sequence types, including 192, 203, and 80 with the *vanB* cassette being much less prevalent overall than the more common *vanA* (27). In fact, only two of our isolates have *vanB*, neither of which were ST117. Similarly, a linear plasmid was identified in ST117 in Switzerland that confers the ability to utilize N-acetyl-galactosamine (GalNAc), a primary constituent of the human gut mucins, as a primary growth substrate (64). But this plasmid was not found in all ST117 – and it was also found in other sequence types. We found no evidence of this plasmid in any of our samples (see Methods).

A recent study also reported that ST117 produces significantly higher proportions of membrane vesicles at subinhibitory concentrations of vancomycin that have been linked to the horizontal transfer of resistance genes like *vanA* or *vanB* (67). We did not test that phenotype in any of our ST117 isolates, but that study reported that ST80 had the same phenotype (67). Finally, a recent study from our group (34) observed that the rise of ST117 and ST80 was correlated with the rise in prevalence of the phenotypically functional bacteriocin bac43 which highlights the need to further study its potential role in transmission and persistence during colonization in health-care environments.

## Conclusion

We found that the population structures of VREfm in the gut and the blood are remarkably similar, that the genetic structure of clones acquired at the hospital is virtually the same as that of those already present in patient arriving to the hospital, and that colonizing strains closely match (genetically) strains obtained from the blood thus highlighting that colonization is a significant infection risk factor. Altogether, this strongly emphasizes the importance of colonization as a risk factor for bloodstream infection and that infection prevention strategies could also focus predominantly in minimizing the chances of VREfm colonizing patients without regard to sequence type. Furthermore, it remains unclear what are the genetic and phenotypic determinants that make ST117 so successful within healthcare environments worldwide. Efforts to identify them can lead to better treatment strategies that minimize the risk of outbreaks, transmission, and further AMR evolution.

## Materials and Methods

### Sampling

Samples were obtained from two sampling strategies at the University of Michigan Hospital. The first sampling strategy was part of a hospital-wide surveillance effort screening patients for the presence of VREfm in the gut between January 1^st^, 2016, and December 31^st^, 2020, through perirectal (PR) swabs from patients admitted into the hospital. Patients were screened every 7 days and positive swabs for VREfm were sent and stored at -80 °C. In the second strategy, samples from all bacteremia (Blood isolates, BL) positive for *Enterococcus sp.* were sent and stored at -80 °C. The BL group includes any species of *Enterococcus* obtained from blood from patients between 2013 and 2020 (from a total of 384 patients and 1925 poly-enterococcal clones). In total we have 748 BL clones from 213 patients, and 74484 PR clones from 39082 patients. FPR of VRE select is 0.96.

### DNA extraction and Library preparation

We extracted DNA from more than 2000 isolates from the PR and BL collections using a Mag Bind Bacterial DNA kit. We streaked frozen isolates on BHI agar and incubated for 48 hours at 35 °C. To complete extraction, we followed the protocol according to manufacturer’s instructions. We then quantified the DNA using a Qubit Fluorometer (ThermoFisher Scientific) to determine high quality and yields. We prepared the library using a Collibri PCR-free ES DNA Library Prep kit for Illumina and submitted the samples for sequencing at the University of Michigan sequencing core. All sequences are publicly available in bioproject PRJNA746456 in NCBI.

### Genome Analysis

We obtained whole-genome sequencing data (WGS) using Illumina HiSeq 125 bp paired-end (68) for a subset of the clones. For analysis we employed an established pipeline encoded in bash and snakemake (69). Briefly, reads were trimmed and filtered for quality using trimmomatic (v0.39) and fastQC (v0.12.1) using a sliding window of 4 and minimum quality score of 25 (4:25) with a minimum read length of 35 (70,71). We then assembled the genomes using unicycler’s (v0.5.0) normal mode (72) and annotated it using bakta’s (v1.8.2) full database (73). From this, we obtained the MLST type for each clone using the mlst software (v 2.23.0-2023-08), which incorporates components of the PubMLST database (74) and obtained antimicrobial resistance associated genes and variants using AMRFinder (v ncbi-amrfinderplus:3.11.18-2023-08-08.2) for *Enterococcus faecium* and *Enterococcus faecalis* organisms (75). We additionally filtered for high-quality genome assemblies (>95% mapping, N50>1000, and at least 30x coverage), resulting in a total of 2061 genomes. All genomes have been previously reported and analyzed for bacteriocin dynamics in VRE in Garretto et al., 2024 (34).

### Plasmid pELF_USZ analysis

To determine the presence and variation of plasmid among our isolates, we aligned all 2061short read sequences to the reference plasmid pELF_USZ (GenBank accession no. OU016038.1) using bwa (v0.7.17-mman5nn). We called variants using FreeBayes (76) with the following parameters: minimum base quality greater than or equal to 10 and minimum alternate fraction less than or equal to 0.5. We then used SnpEff (77) to annotate variants. Then, we identified and annotated insertion elements using panISa (78) and ISFinder (79). Finally, we used SAMtools (80) to identify areas of low read coverage indicating gene deletions and filtered to positions with at least 10% average coverage across the whole plasmid.

### Ethics Statement

This study was approved by the University of Michigan Medicine IRB (IRB #: HUM00102282). There was no direct involvement with patients. The samples obtained were microbiological samples that were obtained for clinical purposes and would have been discarded otherwise.

## Acknowledgements

We would like to thank Kenneta Nunn for her work getting the database setup to complete this endevour.

